# RhoA- and Ran-induced antagonistic forces underlie symmetry breaking and spindle rotation in mouse oocytes

**DOI:** 10.1101/2020.10.20.348045

**Authors:** Benoit Dehapiot, Raphaël Clément, Anne Bourdais, Sébastien Huet, Guillaume Halet

**Affiliations:** Aix Marseille Université, CNRS, IBDM-UMR7288, Turing Center for Living Systems, 13009 Marseille, France; Univ Rennes, CNRS, IGDR - UMR 6290, F-35000 Rennes, France

## Abstract

Mammalian oocyte meiotic divisions are highly asymmetric and produce a large haploid gamete and two small polar bodies. This relies on the ability of the cell to break symmetry and position its spindle close to the cortex before the anaphase occurs. In metaphase II arrested mouse oocytes, the spindle is actively maintained close and parallel to the cortex, until the fertilization triggers the sister chromatids segregation and the rotation of the spindle. The latter must indeed reorient perpendicular to the cortex to enable the cytokinesis ring closure at the base of the polar body. However, the mechanisms underlying symmetry breaking and spindle rotation have remained elusive. In this study, we show that the spindle rotation results from two antagonistic forces. First, an inward contraction of the cytokinesis furrow dependent on RhoA signaling and second, an outward attraction exerted on both lots of chromatids by a RanGTP dependent polarization of the actomyosin cortex. By combining live segmentation and tracking with numerical modelling, we demonstrate that this configuration becomes unstable as the ingression progresses. This leads to spontaneous symmetry breaking, which implies that neither the rotation direction nor the lot of chromatids that eventually gets discarded are biologically predetermined.

## Introduction

Meiosis is the evolutionarily conserved cell division process by which haploid gametes are generated from diploid germ cells. Because meiosis is universally initiated after a pre-meiotic S-phase, two consecutive rounds of cell division - meiosis I and meiosis II - are required to generate gametes with only one copy of each chromosome. In the mammalian female germline, meiotic divisions are highly asymmetric, producing a large oocyte and two smaller polar bodies. This enables the oocyte to conserve most of its cytoplasmic resources (e.g. mRNA, mitochondria, ribosomes) while getting rid of supernumerary genetic material before parental genome fusion. The asymmetry of oocyte meiotic divisions relies on the ability of the cell to break symmetry and position its spindle close to the cortex before the anaphase occurs. The spindle is indeed responsible for setting up the cleavage plane (1) and must therefore be off-centered to allow an uneven partitioning of the gamete. In mitotic cells, spindle positioning is primarily achieved through interactions with the cell cortex, which exerts pulling forces on centrosome-nucleated astral microtubules projecting from the spindle poles (2,3). Intriguingly, oocytes from most species are devoid of centrioles, and therefore lack genuine astral microtubules (4,5). Hence, oocytes must rely on alternate strategies to position their spindle.

Over the last two decades, seminal studies have shed light on how actin filaments (F-actin) play a pivotal role in positioning the spindle during both mouse oocyte meiotic divisions (6–8). During early metaphase I, the spindle assembles in the central region of the oocyte and slowly relocates toward the nearest cortical region (9). The spindle migration depends on a cytoplasmic network of filaments, assembled by the Formin-2 and Spire1/2 nucleators (10–13). This network indeed supports both pushing and pulling forces that are necessary to relocate the spindle (14–18). Once fully migrated, the anaphase I occurs, and half of the homologous chromosomes are discarded into the first polar body. The ovulated oocyte then enters into the prolonged metaphase II arrest, during which the gamete actively maintains its spindle off-centered and parallel to the cortex until fertilization occurs. The metaphase II spindle positioning also depends on the actin cytoskeleton and the emergence of a positive feedback between chromosome positioning and cortical polarization. Indeed, it has been shown that the chromatin generates a gradient of active Ran GTPase (RanGTP) which triggers polarization of the cortex in a dose and distance dependent manner (19,20). This polarization is characterized by an accumulation of actin filaments, referred to as the F-actin cap, encircled by a ring of activated myosin-II (20). Remarkably, the F-actin cap is in turn capable of attracting chromosomes in the cortical vicinity and thus consolidates the polarity by locally increasing the concentration of RanGTP. This can be explained by the ability of the F-actin cap to generate a polarized flow of filaments. Indeed, by doing so, F-actin promotes the emergence of lateral streaming of cytoplasmic materials which converge in the center of the gamete, before pushing back the spindle against the polarized cortex (15,21). A similar directional pushing force was suggested to accelerate spindle migration in meiosis I, once maternal chromosomes are within ~25 μm from the cortex (15). While the molecular details underlying the cortical actomyosin polarization are not yet fully elucidated, we and others have shown that F-actin cap formation requires the polarized activation of the Cdc42 GTPase (Cdc42GTP) and the Arp2/3 complex, downstream of RanGTP (20–26). Inhibiting Ran GTPase, by overexpressing a dominant negative form (RanT24N), leads to a complete loss of the cortical actomyosin polarity and an absence of cytoplasmic streaming (20,21).

The second meiotic division is triggered by sperm entry and leads to the segregation of the sister chromatids (later referred to as the DNA clusters) between the fertilized oocyte and second polar body (PBII). The success of this division relies on the ability of the spindle, which lies parallel to the cortex during metaphase II, to reposition itself perpendicular to the cortex and enable cytokinesis ring closure at the base of the polar body (27). While evidences suggest that spindle rotation is prevented by actin depolymerization, myosin II inhibition and RhoA inactivation(28–30), the underpinnings of this unique symmetry breaking event have long remained elusive. In a recent study, Wang and colleagues provided new insights into this mechanism, showing that rotation is achieved through an asymmetric distribution of forces along the anaphase II spindle (31). It was suggested that spindle rotation arises from cytoplasmic flows of opposite directions at the two ends of the spindle, consecutive to spontaneous symmetry breaking in the distribution of Arp2/3 and myosin-II at the cortex (31).

In the present study, we showed that the rotation results from two antagonistic forces. First, a RhoA dependent ingression of the cytokinetic furrow in the central spindle region and second, a RanGTP dependent attraction of DNA clusters to the polarized cortex. Using numerical modeling, we have demonstrated that the anaphase II spindle configuration becomes unstable as ingression progresses. The slightest initial asymmetry leads to symmetry breaking and spindle rotation. We developed a live 3D segmentation and tracking procedure to confirm these views *in vivo* and showed that, for example, the early positioning of the DNA clusters relative to the cortex biases the orientation of spindle rotation. Overall, this suggests that the rotation of the spindle is the result of a stochastic and spontaneous symmetry breaking process.

## Results

### Stochastic symmetry breaking underlies spindle rotation

To monitor spindle rotation in living oocytes, we injected metaphase II (MII)-arrested oocytes with cRNAs encoding H2B-mCherry and eGFP-MAP4, to respectively label chromosomes and spindle microtubules. After a 3h culture period to allow protein expression, we triggered meiosis II resumption by incubating oocytes in an EtOH complemented culture media (23). This rapidly induced parthenogenetic activation of the gamete, which proceeds to second polar body (PB2) emission. We imaged oocytes at 37°C, using confocal laser scanning microscopy (see selected example in Fig 1A and 1B, S1 and S2 Movies and overall procedure in Methods).

**Fig 1.**
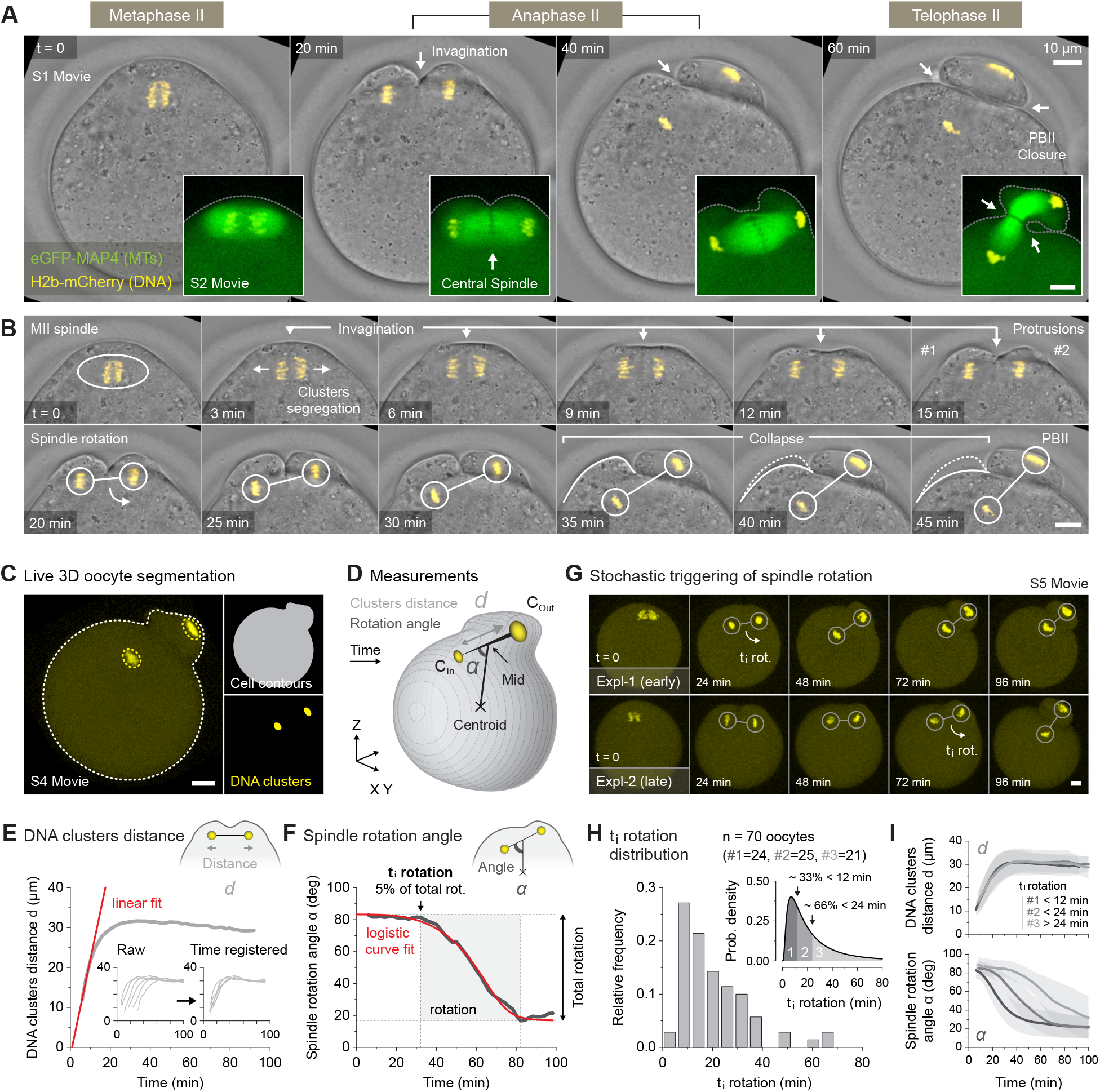
Stochastic symmetry breaking underlies spindle rotation. (A) Live fluorescence and DIC imaging of activated metaphase II oocytes undergoing spindle rotation. The oocytes have been injected either with the H2b-mCherry DNA marker alone (main DIC time series, S1 Movie) or with a combination of H2b-mCherry and the eGFP-MAP4 microtubule marker (inserted spindle time series, S2 Movie). The white arrows show membrane invaginations leading to PBII closure, except in inserted spindle image 20 min where it indicates the central spindle position. (B) High temporal resolution montage of the main DIC time series presented in panel A. The symbols highlight remarkable events occurring during the second meiotic division. (C) Automated 3D segmentation and tracking procedure to monitor the position of the DNA clusters within the shell of the oocyte (see Methods and S4 Movie). (D) Diagram showing the method used to quantify DNA clusters distance *d* and the spindle rotation angle *α*. (E) Variation over time of DNA clusters distance *d* in a selected oocyte (main graph). The red line shows the linear fit used to register in time oocyte population (inserted graphs). (F) Variation over time of spindle rotation angle *α* in a selected oocyte. The red curve shows the logistic fit used to extract rotation parameters. The initial time of rotation (t_i_ rotation) is defined as the time when the rotation reaches 5% of the total fitted rotation. (G) Two examples of H2b-mCherry injected oocytes illustrating the stochastic triggering of spindle rotation (see S5 Movie). The white arrows indicate the approximate start of the rotation. (H) Distribution of the t_i_ rotation for a control population of 70 oocytes gathered from 9 independent experiments (main graph). Probability density function (pdf) of the t_i_ rotation distribution used to determine three equiprobable categories of increasing t_i_ rotation (inserted graph). (I) Averaged variation over time ± s.d. of the DNA clusters distance *d* (top graph) and spindle rotation angle *α* (bottom graph) averaged per category as defined in panel H. All scale bars represent a length of 10 μm.

At anaphase onset, segregated chromatids started to move towards the spindle poles, forming two DNA clusters (see 3 min, Fig 1B). Simultaneously, the cortical region facing the central spindle invaginated and two distinct membrane protrusions assembled above each of the DNA clusters (see 3-15 min, Fig 1B). This is rapidly followed by the rotation of the anaphase II spindle, such that one of the DNA clusters was extruded in the nascent polar body, while the other one remained within the oocyte (see 20-30 min, Fig 1B). Remarkably, the cortical protrusion overlaying the internalized cluster collapsed in the latter stage of the spindle rotation (see 35-45 min, Fig 1B). Eventually, the second meiotic division ended with the cytokinesis ring closure at the base of the polar body (see 60 min, Fig 1A).

Interestingly, using particle image velocimetry (PIV, see Methods), we showed that the triggering of anaphase was also accompanied with dramatic changes in the cytoplasmic streaming pattern (S1A Fig and S3 Movie). Indeed, the post-activation flows were inverted as compared to those observed in metaphase II oocytes (see S1B Fig) (21), with cytoplasmic material flowing away from the polarized domain and moving toward the centre of the gamete (see 15 min S1A Fig). This inversion clearly coincided with the cortical invagination occurring at the central spindle region (S1C and S1D Fig). Remarkably, the collapse of the cortical protrusion (see 40 min S1A Fig) and cytokinesis ring closure (see 40 min S1A Fig) also generated distinct flows in the latter stages of the rotation.

We next devised an automated 3D segmentation and tracking procedure to monitor the position of the DNA clusters within the volume of the oocyte (Fig 1C, S4 Movie and Methods). Using this method, we measured the distance between the two DNA clusters and linearly fitted the early time-points in order to achieve time-registration of all recordings (Fig 1E and Methods). We also measured the spindle rotation angle (as defined in Fig 1D) and used a five-parameter logistic curve fit to extract monotonic smoothed profiles. This allowed us to define the initial time of rotation (hereafter referred to as t_i_ rotation), corresponding to the time when the rotation reaches 5% of the total fitted rotation (Fig 1F and Methods). By looking at various examples (Fig 1G and S5 Movie) or more precisely, at the distribution of the t_i_ rotation (Fig 1H), we observed that the oocytes did not trigger spindle rotation synchronously with regard to the DNA cluster separation (used here as our time reference). This was particularly visible when splitting the oocyte population into three categories, according to their t_i_ rotation (see Fig 1H and 1I). In doing so, one can easily observe that, while distance curves align perfectly, the rotation curves are shifted in time between categories. This high degree of variability indicates that the symmetry breaking process is rather stochastic, and that the dynamics of rotation is not tightly controlled. This led us to further investigate the underlying mechanisms of symmetry breaking.

### The central spindle/RhoA pathway is required for spindle rotation

The RhoA GTPase is a conserved regulator of cytokinesis in animal cells, through the assembly of a contractile cleavage furrow in the cortical region overlying the central spindle (32–34). Accordingly, RhoA activity is required for polar body cytokinesis in mouse oocytes (30,35,36). The canonical RhoA activation pathway involves the guanine nucleotide exchange factor ECT2 (37–39), which is recruited to the central spindle by the so-called centralspindlin complex (MKLP1, MgcRacGAP) (4,40,41). The recruitment of ECT2 relies on the phosphorylation of the MgcRacGAP subunit by Polo-like kinase 1 (PLK1) (42,43). This pathway is recapitulated in the context of the mouse oocyte in the diagram of Fig 2A.

**Fig 2.**
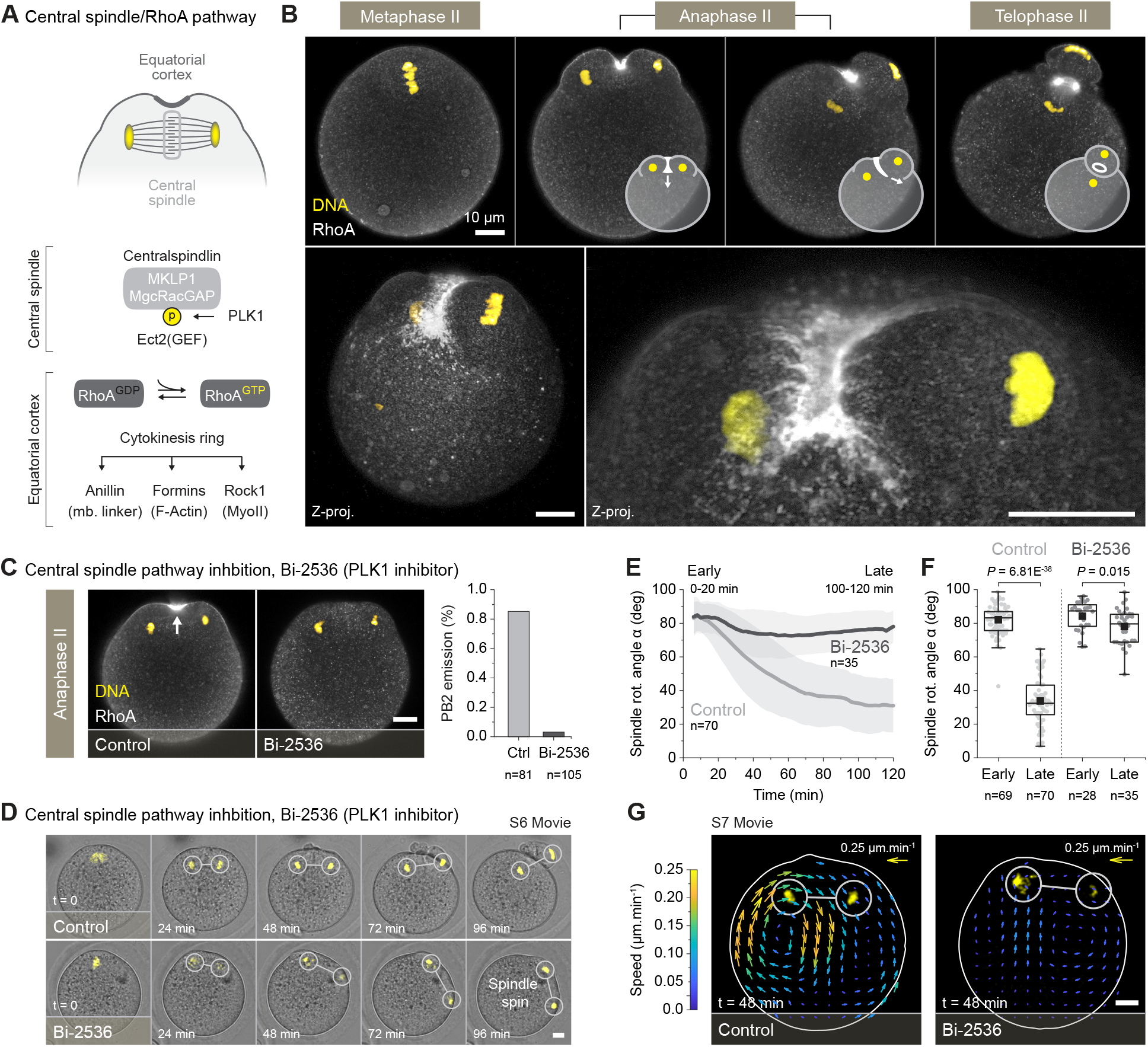
The central spindle/RhoA pathway is required for spindle rotation. (A) Simplified diagram of the central spindle pathway leading to the activation of the RhoA GTPase and the assembly of the cytokinesis ring. (B) Fixed RhoA (immuno-staining) imaging of differentially staged oocytes undergoing spindle rotation. The top panel represents single confocal planes while the bottom panels represent maximum z-projections of multiple confocal planes. The inserted diagrams in the top panel illustrate the progressive closure of the cytokinesis ring (C) Inhibiting the central spindle pathway, using the Bi-2536 (PLK1 inhibitor), prevents the cortical recruitment of RhoA (see white arrow) and the emission of the PBII. The left images show the RhoA localization in fixed oocytes treated with or without the Bi-2536. The right histograms show the PBII emission rate from 81 control and 105 Bi-2536 treated oocytes, both gathered from 4 independent experiments. (D) Live fluorescence and DIC imaging of activated oocytes treated with or without the Bi-2536 (S6 Movie). The oocytes have been injected with the H2b-mCherry DNA marker. (E) Averaged variation over time ± s.d. of the spindle rotation angle *α* in oocytes treated with or without Bi-2536. (F) Box plots: distribution of early (0-20 min) and late (100-120 min) spindle rotation angle *α* in 70 control and 35 Bi-2536 treated oocytes, gathered respectively from 9 and 6 independent experiments. (G) Particle image velocimetry (PIV) measurements, performed on DIC movies, quantifying cytoplasmic flows in a control and a Bi-2536 treated oocyte (S7 Movie). The presented images are extracted from timepoint t = 48 min and show results as a vector field whose color code indicates the speed of tracked particles. The arrow in the top left corner shows the length and the color of 0.25 μm.min^−1^ vector. Box plots in F extend from the first (Q1) to the third (Q3) quartile (where Q3–Q1 is the interquartile range (IQR)); whiskers are Q1 or Q3 ± 1.5 × IQR; horizontal lines represent the median; and black squares represent the mean. Statistics in F were obtained using a two-sided Mann–Whitney test. The exact *p* values are shown directly above the graphs. All scale bars represent a length of 10 μm.

We performed immuno-stainings against RhoA (see Methods) on differentially staged oocytes and succeeded to monitor the GTPase localization throughout the second meiotic division (Fig 2B). Our experiments revealed that, while showing no particular localization during the metaphase II arrest, RhoA strongly accumulated in the cortical region facing the central spindle during anaphase II. This was associated with the recruitment of other classical cytokinesis factor such as ECT2, the scaffolding protein anillin and the Rho-associated protein kinase 1 (Rock-1) (S1 Fig). The RhoA GTPase assembles a hemi-ring invaginating between the two cortical protrusions overlaying the separating DNA clusters (see z-projections, Fig 2B). The hemi-ring remains open until the end of spindle rotation, after which it closes around the central spindle at the base of the polar body (see telophase II, Fig 2B).

We next sought to inhibit the central spindle/RhoA pathway to examine its contribution to spindle rotation. Thus, we treated oocytes with the Bi-2536, a small molecular inhibitor known to specifically target PLK1 in somatic cells (44) and mouse oocytes (45). As expected, PLK1 inhibition prevented the cortical recruitment of RhoA in anaphase, the central cleavage furrow ingression, and resulted in a sharp reduction of the PB2 extrusion rate (controls n=68/81 ~85% PBII, Bi-2536 n=3/105 ~3% PBII) (Fig 2C). When monitored live, the Bi-2536 treated oocytes demonstrated chromatid cluster separation but failed to rotate their spindles. Surprisingly, the latter rather engaged in a spin around the cell periphery before stopping in the latter stages of the division (Fig 2D and S6 Movie). In terms of rotation, this translated in a clear difference between conditions, with spindles in controls undergoing a ~ 50 ° angle variation while the treated ones remained almost parallel to the cortex (mean angle after 120 min, controls 33.7° ± 14.1 S.D., Bi-2536 78.0° ± 10.3 S.D.) (Fig 2E and 2F). Furthermore, the inversion of the cytoplasmic streaming pattern was abolished in the treated oocytes, which suggest that, indeed, the contraction of the cytokinetic furrow is responsible for this phenomenon (Fig 2G and S7 Movie).

Overall, these data indicate that the central spindle/RhoA pathway, which promotes cytokinetic ring assembly and cortical ingression, is required to drive the spindle rotation. Interestingly, upon RhoA inhibition, the spindle still initiates a directional, yet unproductive, spinning movement around the oocyte. This suggests that additional mechanisms may contribute to the symmetry breaking.

### Cortical actomyosin polarization reorganizes during anaphase II

As we have seen previously, the chromosomes carried by the metaphase II generate a gradient of RanGTP that triggers the polarization of the actomyosin cortex in a dose and distance dependent manner (19,20). Remarkably, the polarized cortex is in turn capable of attracting the chromosomes to the cell periphery, allowing the oocyte to keep its spindle off-centered during the metaphase II arrest (21). This ability to modulate the spindle localization encouraged us to investigate whether the DNA-induced polarity pathway still operates during anaphase II and could therefore be involved in the symmetry breaking occurring during spindle rotation.

Consistent with previous observations (18), metaphase II oocytes exhibited a polarized F-actin cap surrounded by a myosin-II ring in the cortical region overlying the spindle (see top panel Fig 3A). In cortical line-scans (see Methods), this took the shape a broad F-actin peak flanked by two smaller myosin-II peaks corresponding to opposite sides of the ring (Fig 3B). This pattern persisted after the anaphase onset but separated in two smaller caps/rings overlaying each of the segregating DNA clusters (see middle panels Fig 3A and 3B). However, in the latter stages of anaphase II, an asymmetry in cortical myosin-II distribution became evident. Indeed, while the ring configuration was maintained in the nascent polar body, the myosin-II invaded the polarized cortex overlying the non-extruded DNA cluster (see bottom panels Fig 3A and 3B). As a result, both the F-actin and myosin-II form a cap in the collapsing protrusion. Note that another cortical actomyosin sub-population emerged in front of the central spindle during anaphase II (arrowheads in middle panels of Fig 3A and 3B). Contrary to the polarized F-actin cap and myosin-II ring, this second sub-population was found to be sensitive to Rock-1 inhibitions (Y-27632), which suggests that it belongs to the cleavage furrow apparatus, and is dependent on the central spindle pathway (see S3 Fig).

**Fig 3.**
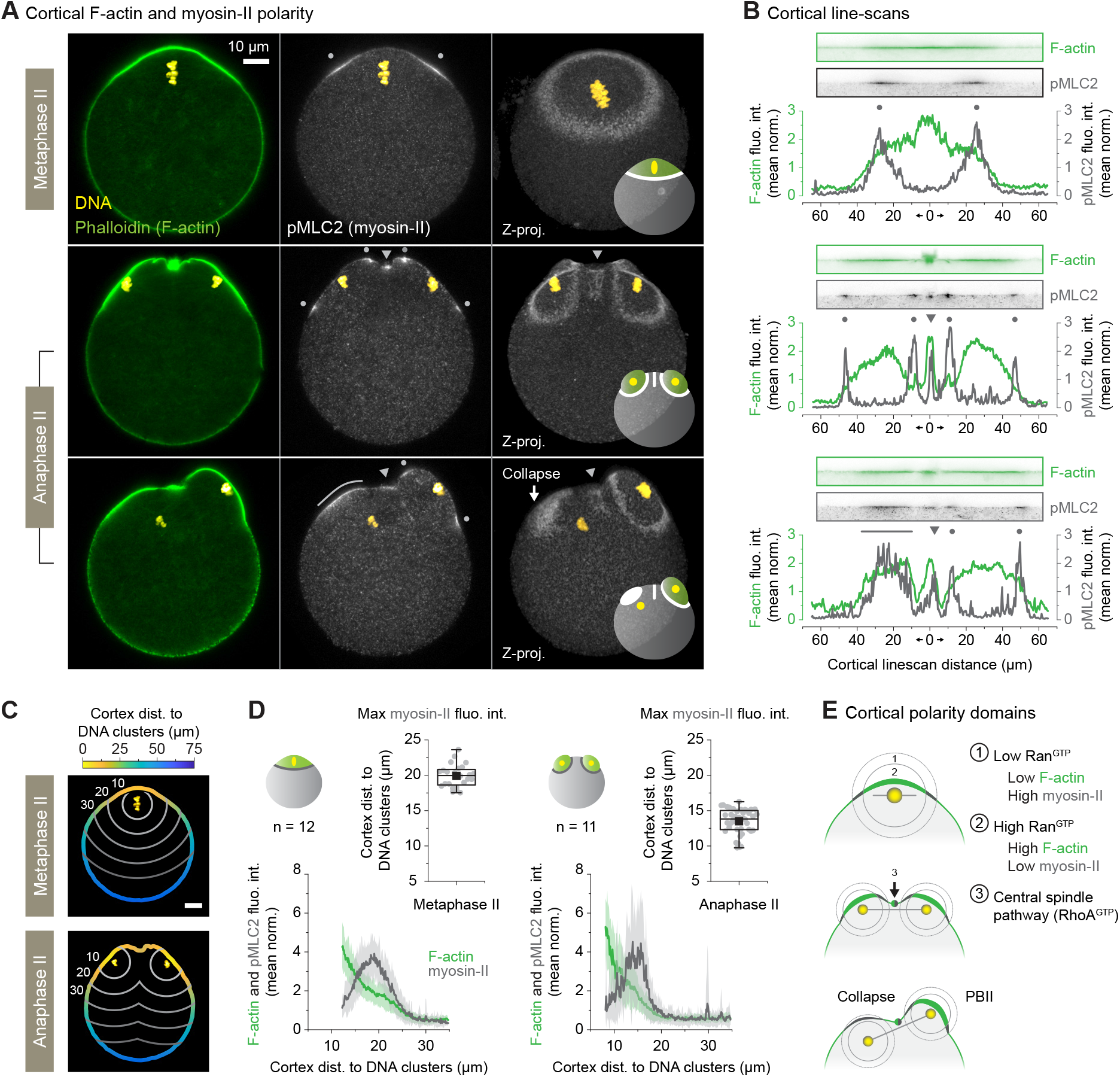
Cortical actomyosin polarization reorganizes during anaphase II (fixed experiments). (A) Fixed F-actin (phalloidin staining) and myosin-II (pMLC2 immuno-staining) imaging of differentially staged oocytes undergoing spindle rotation. The two-left columns of images represent single confocal planes while the right-most column represents maximum z-projections of multiple confocal planes. The grey dots and arrowheads show DNA and the central spindle induced cortical myosin-II populations, respectively. The white arrow shows the collapsing cortical protrusion. The inserted diagrams in the right-most column summarizes the observed cortical actomyosin localization. (B) Cortical fluorescence intensity profiles recapitulating F-actin and myosin-II polarization patterns observed in panel A. The fluorescence intensities are normalized to the mean of each profile. The grey dots and arrowheads show the DNA and the central spindle induced cortical myosin-II populations, respectively (see panel A). Straightened cortical line-scans images used to extract the profiles are shown above the graphs. (C) Diagrams showing the color-coded cortex distance to DNA clusters in both metaphase II (top) and anaphase II oocytes. The grey lines separate oocyte regions equidistant from the DNA clusters. (D) Line graphs: averaged ± s.d. cortical F-actin (phalloidin staining) and myosin-II (pMLC2 immuno-staining) fluorescence intensities as a function of the cortex distance to DNA clusters. The fluorescence intensities are normalized to the mean of each profile. Box plots: distribution of the cortical myosin-II peaks distance to the DNA clusters (see Methods). The quantifications have been performed on 12 metaphase II and 11 anaphase II oocytes, gathered from 2 independent experiments. (E) Diagrams illustrating how the RanGTP gradient, centered on the chromosomes, shapes cortical actomyosin polarization during the second meiotic division. Cortical F-actin accumulates in the proximal part of the gradient (zone 1) while myosin-II concentrates in the periphery (zone 2). Also note the cortical enrichment of actomyosin due to the cytokinesis ring (zone 3). Parameters for box plots in D are as described in Fig 2. All scale bars represent a length of 10 μm.

We next quantified the average cortical F-actin and myosin-II levels as a function of the distance between the cortex and DNA clusters in fixed oocytes (see Methods and Fig 3C and 3D). In both metaphase II and anaphase II oocytes, the cortical F-actin intensity decreased as the distance between the cortex and DNA clusters increased. In contrast, and in line with its ring localization, cortical myosin-II reached its maximum intensity at a certain distance from the DNA clusters (19.9 μm ± 1.6 S.D in metaphase II and 13.5 μm ± 1.8 S.D in anaphase II, see Fig 3D). This indicates that, like in metaphase II, cortical actomyosin polarization is primarily regulated by the proximity of the chromatin in activated oocytes. At close range, RanGTP is high and cortical F-actin is strongly enriched. With distance, F-actin levels decrease and myosin-II assembles as a ring in the distal part of the gradient (top panel, Fig 3E). Half the amount of DNA present in the anaphase II clusters, due to the sister chromatid separation, does not alter this pattern but only reduces the size of the polarization domain (middle panel, Fig 3E). The later recruitment of myosin-II in the collapsing cortical protrusion is consistent with these observations. Indeed, as the spindle rotates, the distance between the cortex and the internalized lot of chromatids increases. As a consequence, only a low dose of RanGTP reaches the cortex at this stage, which promotes the cortical recruitment of myosin-II (bottom panel, Fig 3E).

We pursued our investigation by co-injecting oocytes with the H2b-mCherry DNA marker and a low dose of a cRNA coding for a F-actin probe (eGFP-UtrCH) (46). This allowed us to monitor live cortical F-actin dynamics throughout the division process. Our movies recapitulated what we observed in fixed experiments, with two polarized F-actin caps emerging above the segregating lots of chromatids, one of which becoming the PBII while the other one collapsed in the later stages of division (Fig 4A and S8 Movie). We next wanted to correlate, over time, the cortical level of F-actin with the distance existing between the DNA clusters and the cortex. To do so, we generated color-coded maps of the polarized domain, displaying the spatio-temporal variations of these two quantities (see Methods and Fig 4B). We also determined restricted cortical domains, overlaying the inward and the outward cluster, to average the results and produce 1D profiles (see Methods and Fig 4C). Using both methods we observed a clear correlation between the proximity of the DNA clusters and the level of cortical F-actin polarity. We also noticed, as will be discussed in Fig 6, that the outward cortical domain tended to be more polarized (closer to the DNA cluster and presenting more F-actin) at the onset of spindle rotation (see time = 30 min in Fig 4C).

**Fig 4.**
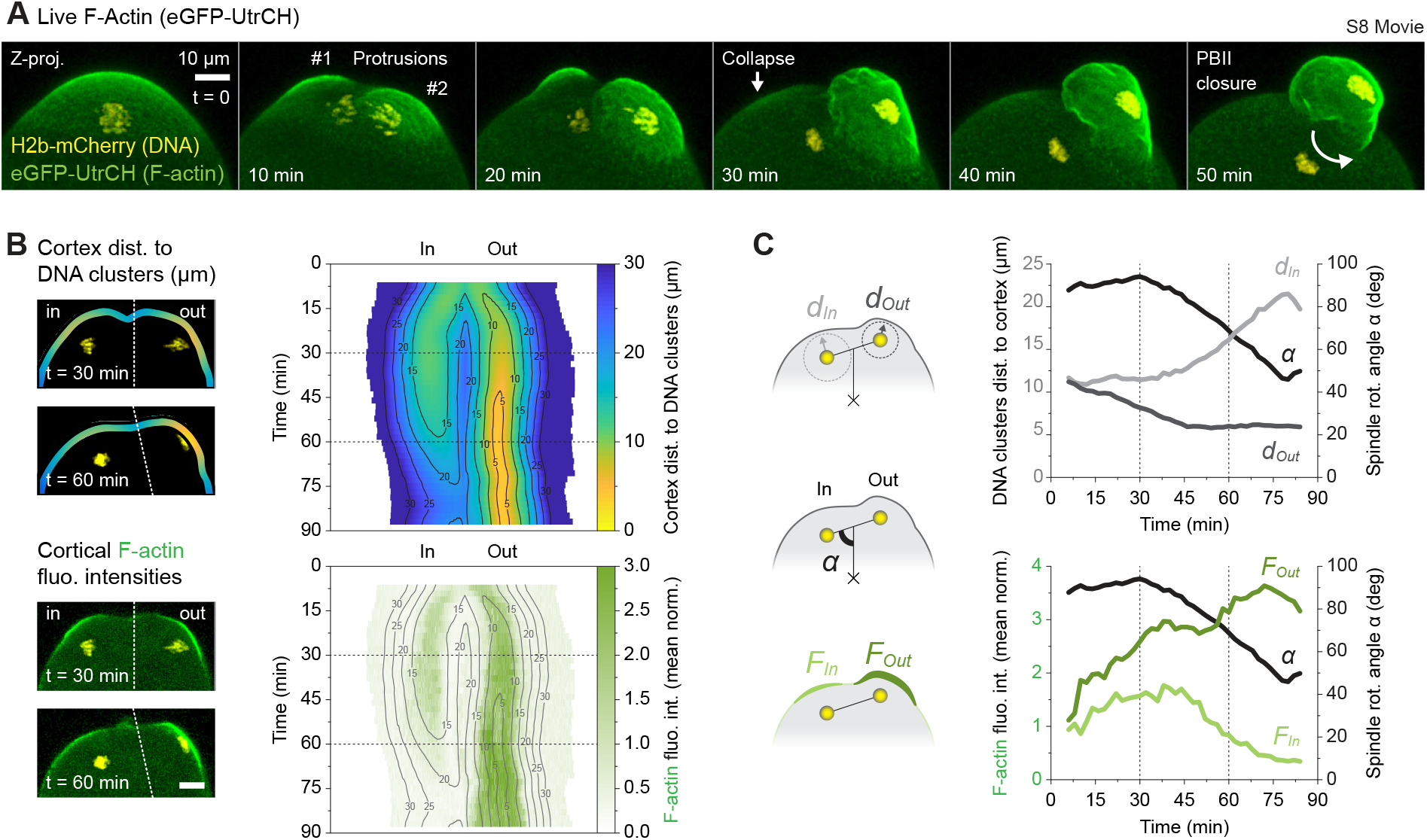
Cortical actomyosin polarization reorganizes during anaphase II (live experiments). (A) Live fluorescence imaging of an activated metaphase II oocyte undergoing spindle rotation. The oocyte has been injected with a combination of the H2b-mCherry DNA marker and the eGFP-UtrCH F-actin marker (S8 Movie). The images represent maximum z-projections of multiple confocal planes and the symbols highlight remarkable events occurring during the second meiotic division. (B) 2D maps of the polarized domain showing variation over time of the cortex distance to DNA clusters (color-coded top panel) and the cortical F-actin levels (bottom panel). The maps are extracted from the images shown on the right. The isodistance lines are used as landmarks to delimit region of the cortex equidistant from the DNA clusters. The dashed lines separate the best focus plane for each DNA cluster. (C) 1D representation of the data shown in panel B (see Methods). The graphs show variation over time of the DNA clusters distance to the cortex (top panel, *d_In_* and *d_Out_*) and the cortical F-actin levels in each polarized domain (see bottom panel, *F_In_* and *F_Out_*). The variation of spindle rotation angle *α* is also represented on both graphs. The diagrams on the left illustrate the measurements described above. All scale bars represent a length of 10 μm.

**Fig 5.**
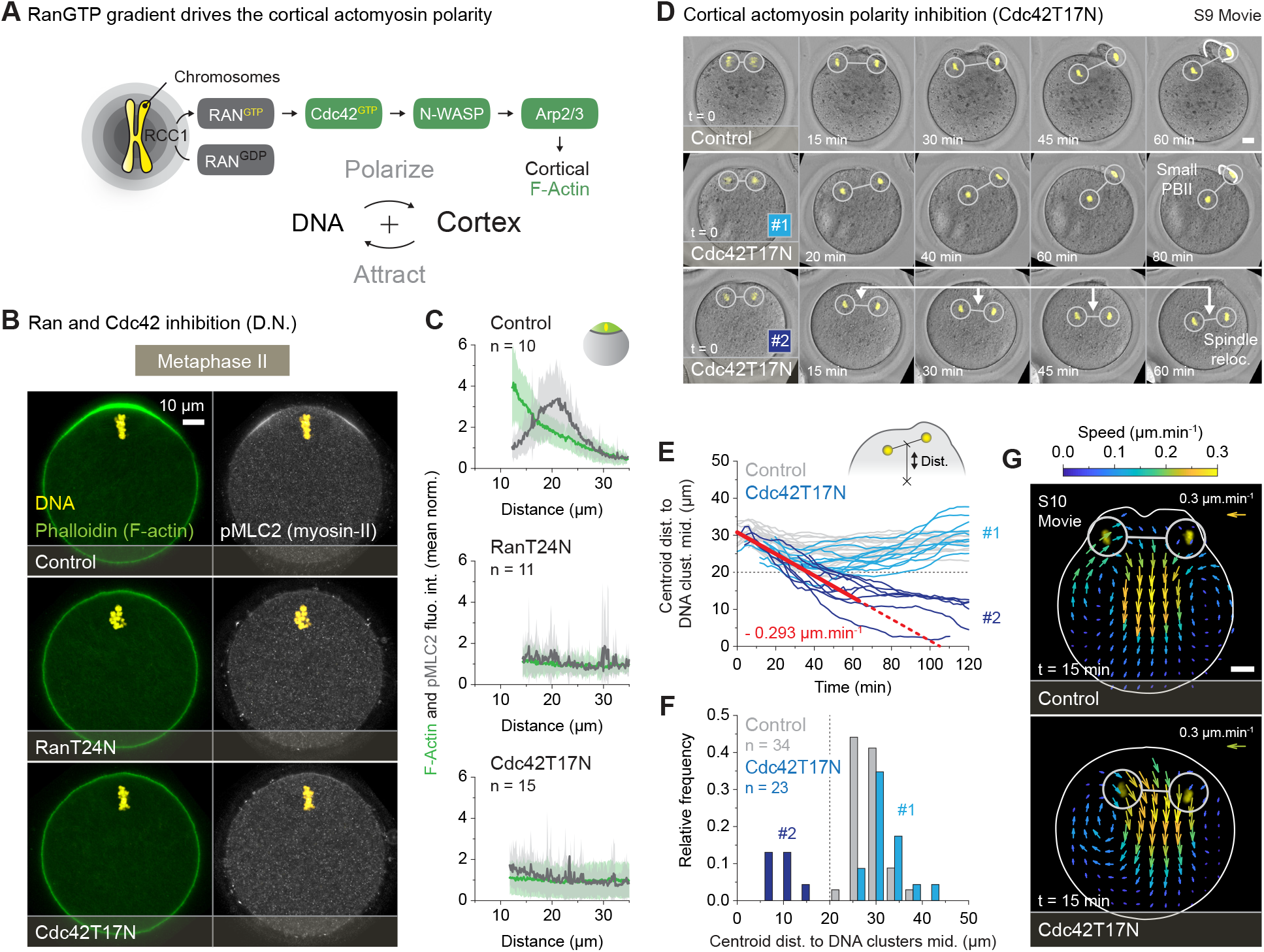
The polarized cortex exerts attraction forces on chromatid clusters. (A) Simplified diagram of the RanGTP gradient, emanating from the chromosomes, and leading to the cortical accumulation of F-actin through the activation of the Cdc42, N-WASP and Arp2/3 pathway. The proximity of the chromosomes promotes cortical F-actin polarity, which in turn attracts the chromosomes to the cortex. (B) Fixed F-actin (phalloidin staining) and myosin-II (pMLC2 immuno-staining) imaging of metaphase II arrested oocytes expressing dominant negative forms of the Ran and Cdc42 GTPase (respectively RanT24N and Cdc42T17N). Control oocytes have been injected with water. (C) Line graphs: averaged ± s.d. cortical F-actin (phalloidin staining) and myosin-II (pMLC2 immuno-staining) fluorescence intensities as a function of the cortex distance to DNA clusters. The fluorescence intensities are normalized to the mean of each profile. Quantifications have been performed on 10 controls, 11 RanT24N and 15 Cdc42T17N injected metaphase II oocytes, gathered from 2 independent experiments. (D) Live fluorescence and DIC imaging of activated oocytes injected with the H2b-mCherry DNA marker alone (control) or a combination of H2b-mCherry and a Cdc42 dominant negative form (Cdc42T17N, group #1 and #2) (S9 Movie). The Cdc42T17N injected oocytes present two characteristic phenotypes, with a first group of gametes extruding a small PBII (as compared to controls) and a second group whose spindle relocates towards the center of the cell. (E) The graph shows variation over time of the distance between the oocyte centroid and the DNA clusters mid-point. The grey profiles represent controls (H2b-mCherry alone) while the light and dark blue profiles represent Cdc42T17N injected oocytes (group #1 and #2, respectively). The red line shows a linear fit, from 0 to 60 min, of the Cdc42T17N group #2. (F) Distribution of the distance between the oocyte centroid and the DNA clusters mid-point after 120 min of recording. Quantifications have been performed on 34 control and 23 Cdc42T17N injected oocytes, gathered respectively from 9 and 3 independent experiments. (G) Particle image velocimetry (PIV) measurements, performed on DIC movies, quantifying cytoplasmic flows in a control and a Cdc42T17N injected oocyte (S10 Movie). The presented images are extracted from timepoint t = 15 min and show results as a vector field whose color code indicates the speed of tracked particles. The arrow in the top left corner shows the length and the color of 0.3 μm.min^−1^ vector. All scale bars represent a length of 10 μm.

**Fig 6.**
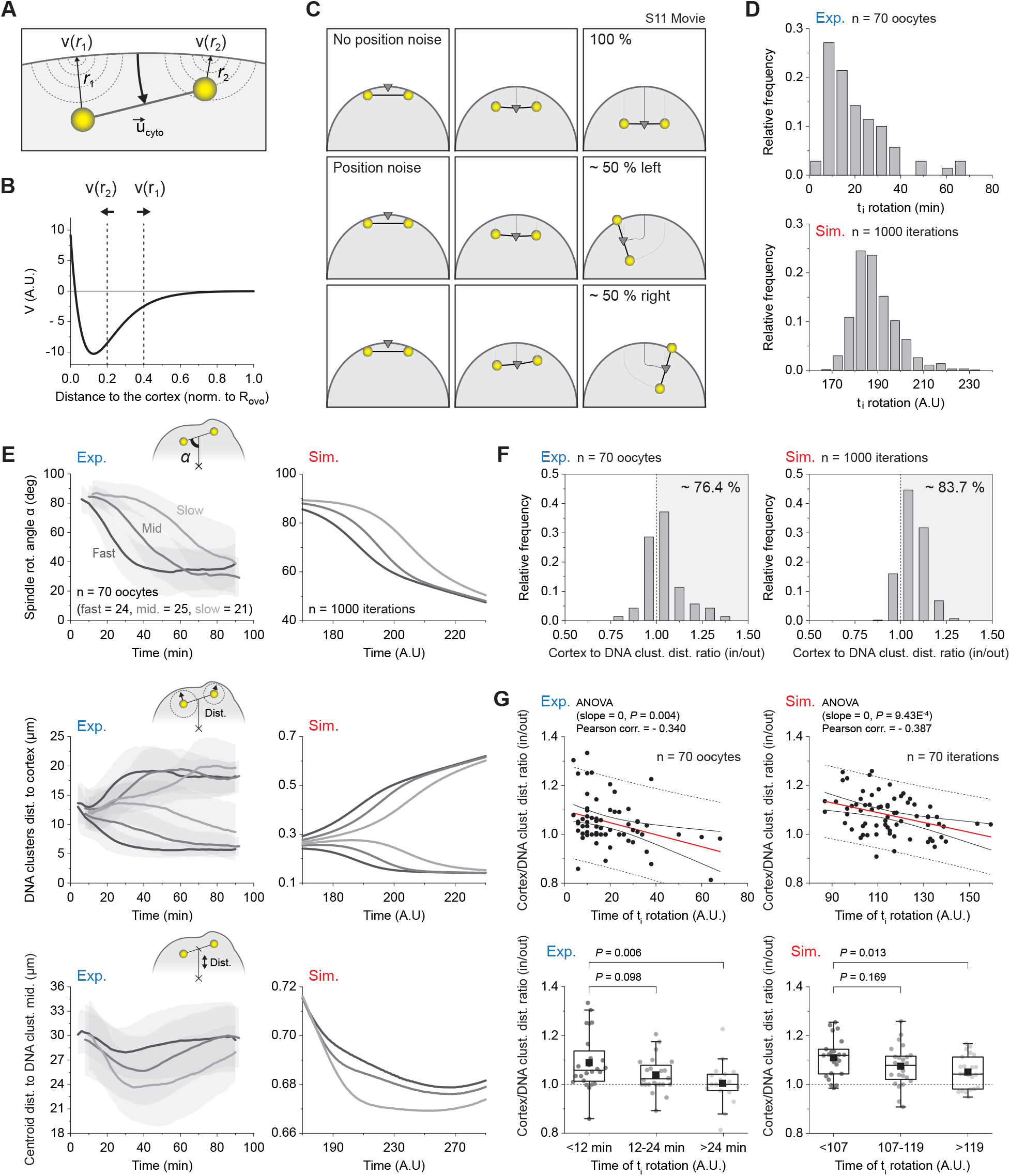
Antagonistic forces yield symmetry breaking and spindle rotation. (A) Diagram recapitulating the main elements used in the numerical model. A constant inward velocity, 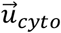, has been applied to the spindle mid-point to recapitulate the cytokinetic furrow ingression. The attraction of the DNA clusters for the cortex has been modelled through an effective potential, *V*(*r*). The strength of this potential decays exponentially as the DNA clusters distance to the cortex increases (see *r*_1_ and *r*_2_). See Methods for further explanation. (B) Shape of the effective potential, *V*(*r*), modelling the attraction of the DNA clusters for the cortex. (C) Time-lapse sequences of typical simulations with (top row) or without (middle and bottom rows) Gaussian noise added to the DNA cluster position (S11 Movie). (D) Experimental (top) and simulated (bottom) distributions of the initial rotation times (t_i_ rotation). (E) Experimental (left column) and simulated (right column) variation over time ± s.d. of the spindle rotation angle *α* (top row), the DNA clusters distance to the cortex (middle row) and the distance between the oocyte centroid and the DNA clusters mid-point (bottom row). For each graph, the oocyte population has been separated into 3 categories of increasing t_i_ rotation (see Fig 1H). (F) Experimental (left) and simulated (right) distributions of the averaged DNA clusters distance to cortex ratio (in/out) in the first 10 minutes of the recordings. The values above one (see grey background) indicate that the externalized cluster (out) was closer to the cortex before the rotation occurs. (G) Experimental (left column) and simulated (right column) averaged DNA clusters distance to cortex ratio (in/out) versus t_i_ rotation, shown as scatter plots (top row) or binned box plots (bottom row). The red line in the scatter plots represents a linear fit of the point cloud. The plain black lines delimit the 95 % confidence band while the dashed lines delimit the 95 % prediction band. The experimental results in D, E, F and G have been obtain from a population of 70 control oocytes, gathered from 9 independent experiments. The simulated results in D, E and F have been obtain from 1000 simulations, while in G this number have been reduce to 70 to match the size of the experimental data. Parameters for box plots in G are as described in Fig 2. Statistics in G (top row) were obtained using a one-way ANOVA to test the influence of the averaged DNA clusters distance to cortex ratio (in/out) over the t_i_ rotation. Statistics in G (bottom row) were obtained using a two-sided Mann–Whitney test. In both cases the exact *p* values are shown directly above the graphs.

Altogether, the above findings demonstrate that the DNA-induced polarity pathway, operating during metaphase II, is still at play after the anaphase onset. The difference here is that the polarized cortex divides in two smaller domains, above each of the segregating DNA clusters. The polarization intensity of the sub-domains evolves during the rotation and depends on the proximity of the chromatin.

### The polarized cortex exerts attraction forces on chromatids clusters

To further examine how the cortical actomyosin may influence spindle rotation, we next sought to inhibit oocyte cortical polarization by targeting the Ran GTPase signaling pathway (see diagram Fig 5A). To do this, we either targeted Ran itself or its downstream effector Cdc42 by using dominant negative overexpressions (respectively RanT24N (47) and Cdc42T17N (48)). Cdc42 promotes cortical F-actin polarization in metaphase II oocytes through the activation of N-WASP and the Arp2/3 complex (Fig 5A) (21,23,25). We found that, when overexpressed for 3 to 4 hours, both dominant negative forms resulted in a complete loss of the F-actin and myosin-II cortical polarity in metaphase II oocytes (see Fig 5B and the flattening of the polarization curves in Fig 5C).

Using the same strategy, we next examined the effects of Cdc42 inhibition in activated oocytes. We did not consider inhibiting the upstream Ran GTPase, as RanGTP plays an additional role in maintaining spindle integrity (19,49). Thus, we recorded confocal time-series of parthenogenetically activated oocytes expressing Cdc42T17N and H2b-mCherry, and we observed two distinct phenotypes (Fig 5D and S9 Movie). In the first group of oocytes (n = 16/23 ~70%), the anaphase spindle remained off-centered (spindle-centroid distance > 20 μm) and a PB2 protruded, though smaller in size than controls. In the second group (n = 7/23 ~30%), the anaphase spindle relocated substantially toward the oocyte center (spindle-centroid distance < 20 μm), and PB2 was not emitted (see Methods and Fig 5E and 5F). As we have previously shown that overexpressing Cdc42T17N alone inhibits PB2 emission in a large majority of activated oocytes (23), we considered the small polar body phenotype as an incomplete Cdc42 inhibition. Indeed, it is likely that co-injecting Cdc42T17N with H2b-mCherry resulted in a decreased expression of the dominant negative form and thus of its ability to compete with the endogenous GTPase. We therefore assumed the spindle relocation as the real effect of a successful Cdc42 inhibition.

We further noted Cdc42T17N injected oocytes were still undergoing cell-wide cytoplasmic streaming after the anaphase onset (Fig 5G and S10 Movie). This observation prompted us to investigate whether the inversion of the flows could be responsible for the spindle relocation upon Cdc42 inhibition. Indeed, if this is the case, this would indicate that Cdc42 activity is necessary to counteract the flow and maintain the spindle off-centered during division. To answer this question, we measured the speed of cytoplasmic streaming using PIV and compared it to the speed at which the meiotic spindle relocated in Cdc42T17N injected oocytes (see linear fit in Fig 5E). In both cases, we measured a speed of about 0.3 μm.min^−1^, showing that the spindle and the cytoplasmic materials were moving simultaneously towards the center of the gamete. This suggests that indeed the post-activation flows exert a pushing force on the meiotic spindle that cannot be resisted if Cdc42 is inhibited. Since it has been reported that the active form of the GTPase only accumulates in the cortical protrusions during the anaphase II (23), it can be assumed that Cdc42 counteracts the flow by its ability to assemble the polarized actomyosin cortex, which is known to exert attractive forces on the chromosomes (15,20,21). Other scenarios, such as a role of Cdc42 in anchoring the spindle to the cortex (see Fig 1A), seem less likely given GTPase localization.

Our Cdc42 loss of function experiments showed that, as in metaphase II, the polarized domains exert attractive forces on the chromosomes. However, due to the spindle relocation phenotype, observed upon Cdc42 inhibition, we were not able to make conclusions about the potential involvement of the polarized cortex in the symmetry breaking. We therefore conducted a last series of experiments, mixing numerical modeling and correlative analyses, in an attempt to provide new elements to answer this question.

### Antagonistic forces yield symmetry breaking and spindle rotation

Based on our findings, we considered that the anaphase II spindle is mainly subject to two antagonistic forces: 1) the inward contraction of the cytokinesis ring occurring in the central-spindle region and 2) the outward attraction of the DNA clusters to their respective polarized cortical domain. To translate this in mathematical terms, we devised a simplified model in which a semi-flexible spindle connects two DNA clusters encapsulated in a circular oocyte (see Methods and Fig 6A). To simulate the cytokinetic furrow ingression we prescribed a constant inward velocity to the spindle mid-point, oriented perpendicular to the spindle axis. On the other hand, we modelled the attraction of DNA clusters using an effective potential, akin to a soft-core potential (see the potential profile in Fig 6B), accounting for a feedback such that the attraction of the clusters increases as their distance to the cortex decreases.

As the ingression progresses, the attraction domains and the spindle mid-point are no longer aligned, which prevents DNA clusters to remain both apposed to the cortex. In the absence of numerical noise, the simulation remained perfectly symmetric and the spindle was simply pushed, parallel to the cortex, inside the oocyte (top panel Fig 6C and S11 Movie) However, this configuration was actually unstable, and the addition of a small Gaussian noise to the clusters position (see Methods) was enough to break the symmetry and systematically trigger the spindle rotation (bottom panels Fig 6C and S11 Movie). Rotation resulted in a more energetically favourable configuration, with one of the DNA clusters returning to its potential well, while the other eventually escaped from it (see arrows in Fig 6B).

We next analysed the dynamics of symmetry breaking in simulations and observed that, as in experiments, the distribution of the initial time of rotation (t_i_ rotation, defined as in Fig 1F) was spread out over time and was skewed to the right (Fig 6D). We used this temporal difference to categorize our real and simulated oocyte populations into three categories, according to how soon they break symmetry. This allowed us to evaluate our model and compare how the time at which the rotation starts, influenced relevant geometric quantities such as the spindle rotation angle *α*, the DNA clusters’ distance to cortex and the depth of ingression in real and simulated oocytes (Fig 6E). We found that our model nicely recapitulated the dynamics of symmetry breaking with delayed oocytes (top panel) maintaining their clusters at similar distance from the cortex for a longer period (middle panel) and undergoing a deeper ingression of the cytokinetic furrow (bottom panel).

In the light of this model, and our observations (see Fig 4C), a straightforward prediction was that an initial asymmetry in the DNA clusters’ position should bias the direction of rotation. To verify this hypothesis, we measured the distance to the cortex of each of the two DNA clusters (in and out) and looked at the distribution of the averaged DNA cluster distance to cortex ratio (in/out) in the first 10 minutes of the recordings (Fig 6F). We observed that, in both simulations and experiments, the distributions were biased to the right, indicating that the externalized cluster (out) tended to be closer to the cortex before the rotation occurs (experiments 76.4 %; simulations 83.7% right bias). In addition, we also predicted that a higher initial asymmetry should lead to an earlier start of rotation. We therefore plotted the ratio (as defined above) against the rotation start times and found that these two quantities were negatively correlated in both simulations and experiments (Fig 6G). This confirmed that the delayed oocytes were initially more symmetric.

Finally, this led us to question whether this simple instability mechanism could also recapitulate the right-tailed distribution of the t_i_ rotation (Fig 6D). Running a batch of simulations with Gaussian noise, we found that the distribution of t_i_ rotation was not Gaussian but skewed to the right, as observed experimentally (Fig 6D). To make further sense of this non-trivial observation, we devised a simplified analytic model of symmetry breaking, in which we calculated the time required to reach an arbitrary symmetry breaking point from an initial configuration that slightly deviated from the unstable equilibrium. We could then calculate the distribution of the symmetry breaking time when the initial offset had a Gaussian distribution. We found that the symmetry breaking time had a right-tail distribution, which was a direct consequence of the non-linear dependency of the rotation time on the initial offset (see Methods and S4 Fig).

Overall, this model shows that rotation occurs due to the instability of the parallel configuration as the inward ingression of the cytokinetic furrow progresses. A minimal set of ingredients, namely the inward contraction and the lateral outward attractions, are sufficient to nicely recapitulate the dynamic and stochastic aspects of spindle rotation.

## Discussion

Overall, this study reveals that the DNA-induced polarity pathway operating in metaphase II is still active after anaphase onset and underlies the symmetry breaking occurring during spindle rotation. The difference here is that the polarized cortex splits into two smaller-sized domains overlying each of the segregating DNA clusters. Like in metaphase II, the polarized cortex exerts attractive forces on the clusters which, coupled to the contraction of the cytokinesis ring in the central spindle region, leads to an unstable configuration resulting in spindle rotation. Using theoretical modelling, we showed that this instability stems from the geometric incompatibility of maintaining both DNA clusters at the cortex while pushing the spindle midpoint inward upon cytokinetic ingression. Consequently, any initial asymmetry in the system is sufficient to break symmetry and trigger spindle rotation. Our *in vivo* measurements confirmed these views, as the initial relative distance of the DNA clusters to the cortex before rotation proved to be a good predictive marker of its orientation.

In the absence of clear data on the precise origin and nature of the attractive forces acting on the DNA clusters, we used an effective attraction potential in our model. This allowed us to remain general and only speculate about the involved forces. Hence, our model is centred on a minimal set of ingredients, and yet recapitulates the non-trivial dynamics and stochasticity observed experimentally. In agreement with Wang and colleagues (31), we are inclined to believe that cytoplasmic streaming provides the necessary forces for spindle rotation. However, from our perspective, the unilateral contraction of the RanGTP-induced myosin-II ring (see Fig 3), may not be responsible for the inversion of the flow, described by these authors as essential for spindle rotation. Indeed, our PIV analysis showed that the inversion occurs early in anaphase II, at a time when none of the cortical protrusions has yet collapsed (S1 Fig). This indicates that the myosin-II still forms a ring in both protrusions and that the contraction of the latter cannot be held responsible for the inversion. Furthermore, Cdc42 inactivated oocytes still undergo reverse cytoplasmic streaming despite the loss of both the F-actin cap and the myosin-II ring (see Fig 5). In our view, the inversion of the flow is most likely driven by the contracting cytokinetic furrow, consistent with the strong time correlation of these two phenomena (S1 Fig).

If the myosin-II ring is not responsible for the early inversion of the flow we can therefore speculate its other potential functions. First, we cannot exclude that the contraction of the ring contributes to cytoplasmic streaming to some extent, as well as spindle rotation in the later stages of anaphase II. It is also possible that its contraction drives the rapid collapse of the internalizing cortical protrusion, thereby preventing it from interfering with the PBII extrusion. Finally, and more generally, the DNA-induced myosin-II ring may be a means for the cell to spatially constrain the F-actin cap and thus better control the attractive forces generated by the polarized cortex. In any case, it will be difficult to draw conclusions about the role of the myosin-II ring and its subsequent contraction without identifying a specific way to inhibit it during the spindle rotation. Unfortunately, broad inhibitors such as blebbistatin and ML-7, which effectively prevent myosin-II ring assembly in MII oocytes, were also shown to prevent cytokinesis ring contraction (20,29). On the other hand, we show that the Rock-1 inhibitor, Y-27632, specifically inhibits myosin-II in the cytokinesis furrow, showing a specificity for the cytokinesis ring myosin-II subpopulation without affecting the polarized rings (S3 Fig). This raises the idea that perhaps another kinase, acting downstream of RanGTP, might specifically promote the assembly of the polarized myosin-II rings.

To conclude, as it is often the case in living systems (50), the symmetry breaking process underlying spindle rotation arises from the ability of the oocyte to detect and amplify an initial asymmetry. This is somehow similar to what happen in meiosis I, during which the metaphase I spindle builds upon a slight off-centering to determine the direction of its migration (9). During meiosis II, our results ultimately suggest that the direction of the rotation, and thus which set of chromatids is eventually discarded, is not biologically predetermined but rather results from a stochastic, spontaneous symmetry breaking process.

## Acknowledgements

We are grateful to the staff of the ARCHE-Biosit animal facility and MRIC-Biosit microscopy facility for technical assistance and expert advice. This work was supported by a CNRS ATIP fellowship to GH and the Ligue contre le cancer Grand Ouest (committees 35 and 56). G.H. wishes to thank the Society for Reproduction and Fertility (SRF) for the award of an Academic Scholarship. B.D. received a PhD scholarship from the French Ministry of Research and Higher Education, and complementary funding from the Fondation pour la Recherche Médicale. The authors declare no competing financial interests.

---

**S1 Fig.**
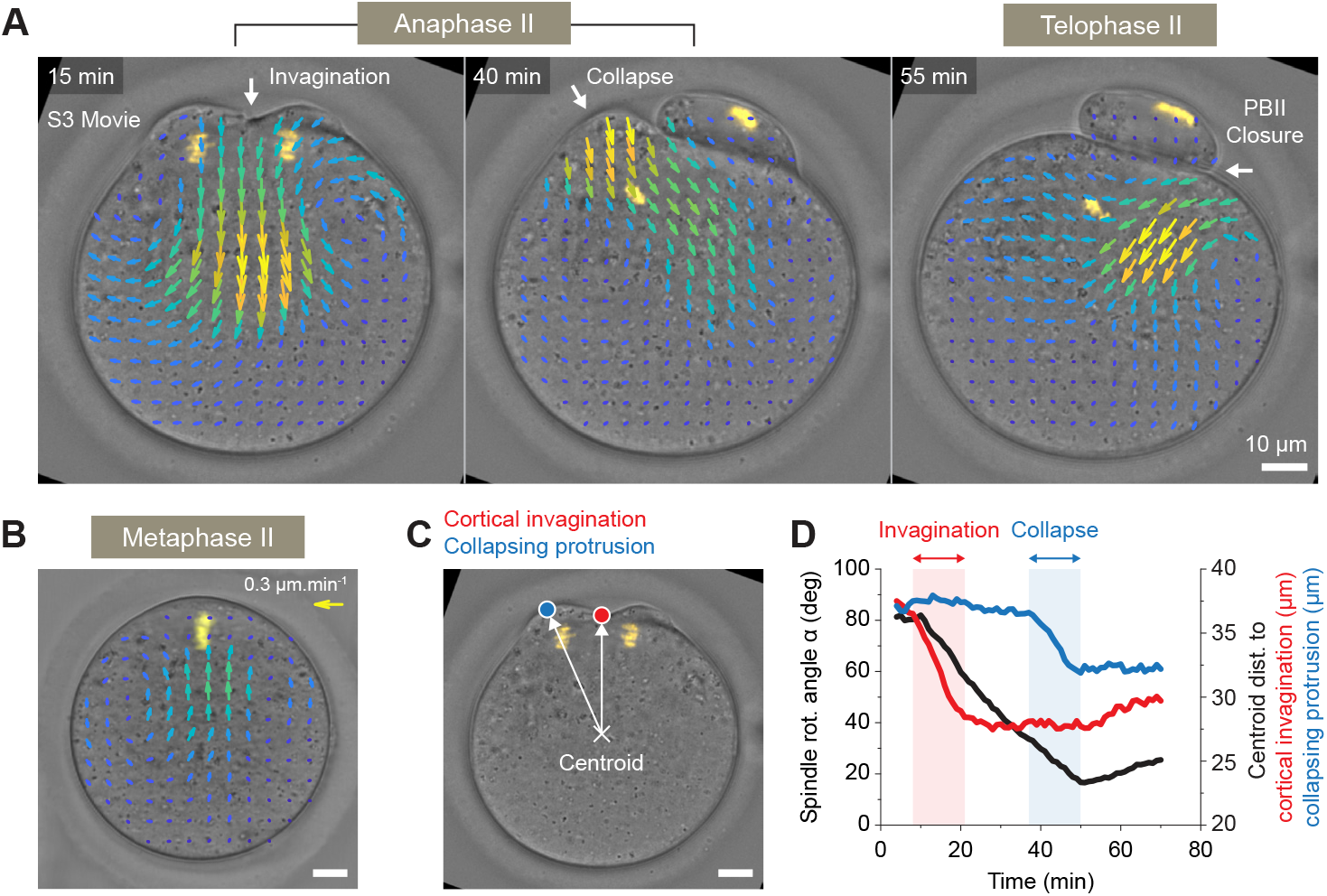
Cytoplasmic streaming during spindle rotation. (A) Particle image velocimetry (PIV) measurements, performed on a DIC movie, quantifying cytoplasmic flows in an activated metaphase II oocyte (S4 Movie). The presented images show the reversion of flow induced by the cytokinetic furrow ingression (t = 15 min), the flow induced by the collapsing protrusion (t = 40 min) and the flow induced by the closure of the cytokinesis ring (t = 55 min). The vector field color code indicates the speed of tracked particles. (B) PIV measurements, performed on a DIC movie, quantifying cytoplasmic flows in a metaphase II arrested oocyte. Contrary to activated oocyte, the cytoplasmic streaming is directed towards the spindle and the polarized cortex. The vector field color code indicates the speed of tracked particles. The arrow in the top left corner shows the length and the color of 0.3 μm.min^−1^ vector. (C) Diagram illustrating the centroid distance to the cortical invagination (red dot) or the collapsing protrusion (blue dot). (D) The graph shows variation over time of the spindle rotation angle *α* (black curve) and the centroid distance to the cortical invagination (red curve) or the collapsing protrusion (blue curve). All scale bars represent a length of 10 μm.

**S2 Fig.**
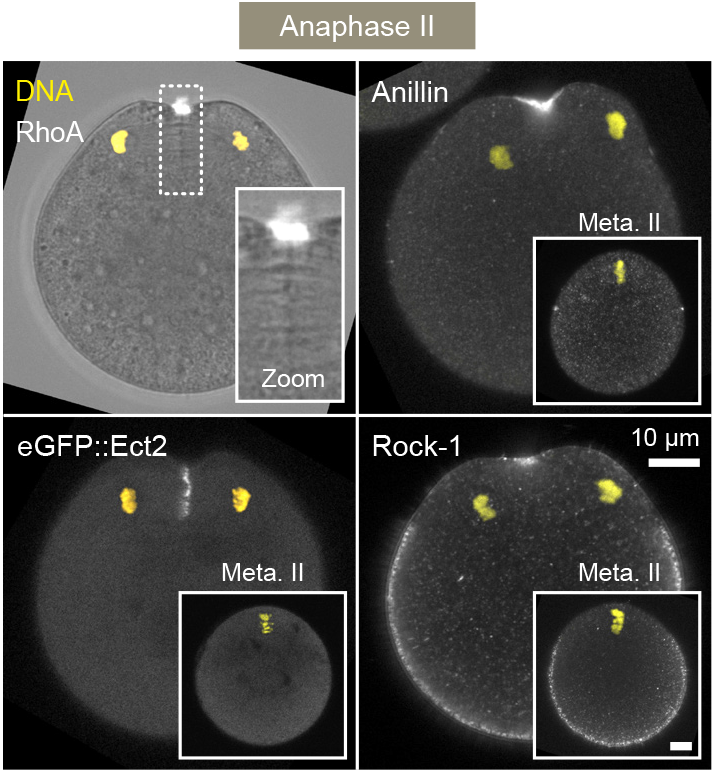
Recruitment of various classical cytokinesis factors during anaphase II. Fixed RhoA, anillin, Rock-1 (immuno-staining) and live eGFP-Ect2 imaging of metaphase II (insets) and activated oocytes (main images). The scale bar represent a length of 10 μm.

**S3 Fig.**
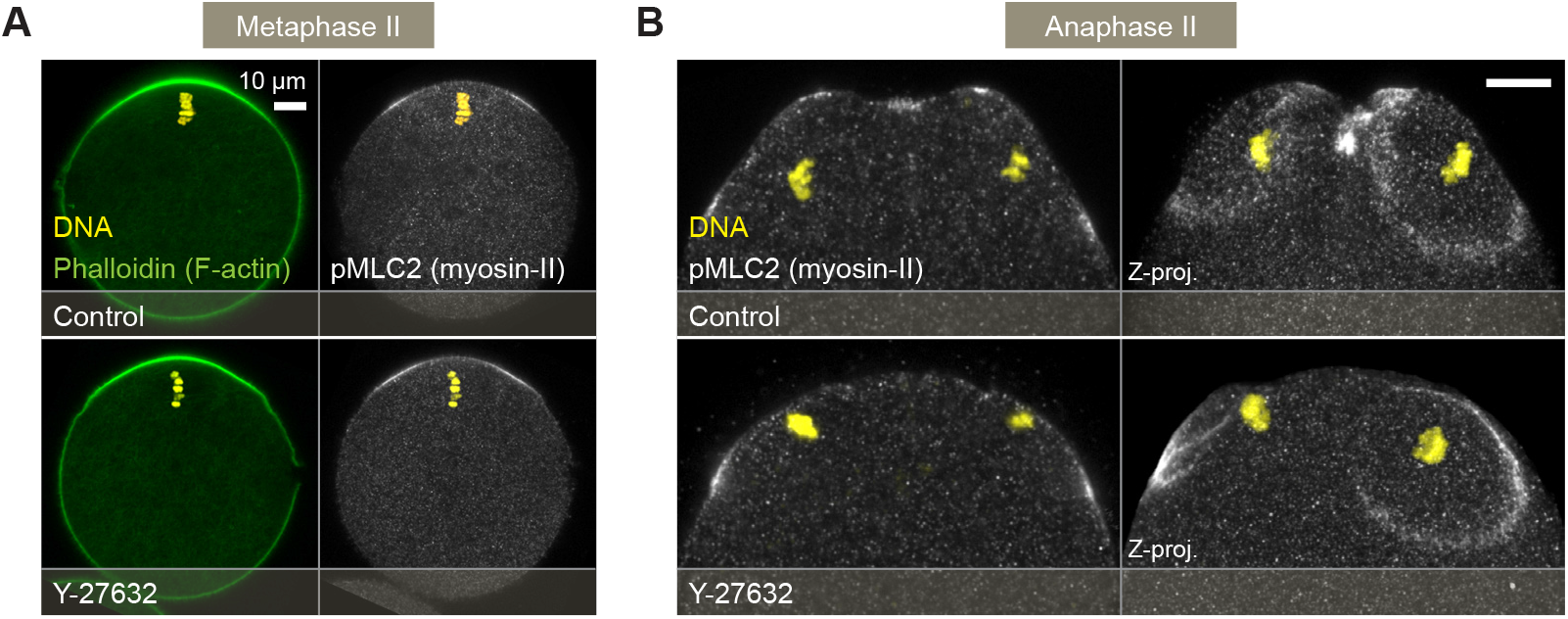
Rock-1 inhibition revealed differentially regulated cortical myosin-II sub-populations. (A) Fixed F-actin (phalloidin staining) and myosin-II (pMLC2 immuno-staining) imaging of metaphase II arrested oocytes treated with or without Y-27632. (B) Fixed myosin-II (pMLC2 immuno-staining) imaging of activated metaphase II oocytes treated with or without Y-27632. The left column shows single confocal planes while the right column shows maximum z-projections of multiple confocal planes. All scale bars represent a length of 10 μm.

**S4 Fig.**
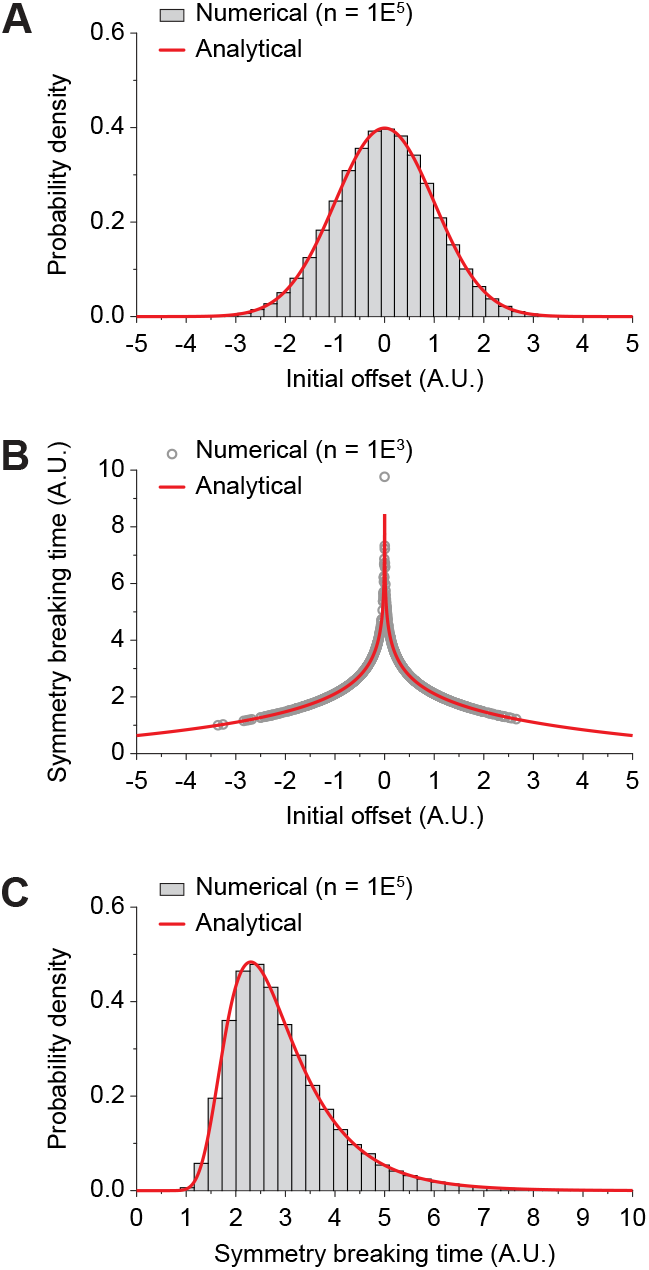
Simplified analytic model of symmetry breaking. (A) Gaussian distribution of initial deviation from the unstable configuration. The red solid line shows the analytical distribution. (B) Distribution of the resulting symmetry breaking time, required to reach an arbitrary distance from the unstable configuration. The red solid line shows the analytical distribution. (C) Symmetry breaking time v.s. initial offset. The red solid line shows the analytical expression.

## Movie legends

**S1 Movie. Stochastic symmetry breaking underlies spindle rotation (DIC).**

Live fluorescence and DIC imaging of an activated metaphase II oocyte undergoing spindle rotation. The oocyte have been injected with the H2b-mCherry DNA marker. The images were acquired every 1 min with a 63x objective and represent a single confocal Z-plane. Scale bar = 10 μm.

**S2 Movie. Stochastic symmetry breaking underlies spindle rotation (MTs).**

Live fluorescence imaging of an activated metaphase II oocyte undergoing spindle rotation. The oocyte have been injected with a combination of the H2b-mCherry DNA marker and the eGFP-MAP4 microtubule marker. Images were acquired every 2 min with a 20x objective and represent a maximum projection of 20 confocal Z-planes. Scale bar = 10 μm.

**S3 Movie. Cytoplasmic streaming during spindle rotation (PIV).**

Live fluorescence and DIC imaging of an activated metaphase II oocyte undergoing spindle rotation. The oocyte have been injected with the H2b-mCherry DNA marker. Images were acquired every 1 min with a 63x objective and represent a single confocal Z-plane. The PIV vector field enables to visualize cytoplasmic streaming and the color code indicates the speed of tracked particles. Scale bar = 10 μm.

**S4 Movie. Automated 3D segmentation and tracking procedure.**

Automated 3D segmentation and tracking procedure to monitor the position of the DNA clusters within the shell of oocyte. The top panel shows tracked DNA clusters from 3 differentially oriented oocytes, while the bottom panel shows a 3D reconstruction of the same oocyte volumes. Images were acquired every 2 min with a 20x objective and represent a maximum projection of 20 confocal Z-planes.

**S5 Movie. Stochastic triggering of spindle rotation.**

Live fluorescence imaging of two activated metaphase II oocytes showing a clear difference in their rotation initiation time. The oocyte have been injected with the H2b-mCherry DNA marker. Images were acquired every 2 min with a 20x objective and represent a maximum projection of 20 confocal Z-planes. Scale bar = 10 μm.

**S6 Movie. Central spindle/RhoA pathway inhibition (PLK1 inhibitor, Bi-2536).**

Live fluorescence and DIC imaging of activated oocytes treated with or without the PLK1 inhibitor, Bi-2536. The oocyte have been injected with the H2b-mCherry DNA marker. Images were acquired every 2 min with a 20x objective and represent a maximum projection of 20 confocal Z-planes (H2b-mCherry) or a single confocal Z-plane (DIC). Scale bar = 10 μm.

**S7 Movie. Central spindle/RhoA pathway inhibition (PLK1 inhibitor, Bi-2536) (PIV).**

Live fluorescence and DIC imaging of activated oocytes treated with or without the PLK1 inhibitor, Bi-2536. The oocyte have been injected with the H2b-mCherry DNA marker. Images were acquired every 2 min with a 20x objective and represent a maximum projection of 20 confocal Z-planes (H2b-mCherry) or a single confocal Z-plane (DIC). The PIV vector field enables to visualize cytoplasmic streaming and the color code indicates the speed of tracked particles. Scale bar = 10 μm.

**S8 Movie. Cortical F-actin polarization reorganizes during anaphase II.**

Live fluorescence imaging of an activated metaphase II oocyte undergoing spindle rotation. The oocyte has been injected with a combination of the H2b-mCherry DNA marker and the eGFP-UtrCH F-actin marker. Images were acquired every 2 min with a 20x objective and represent a maximum projection of 20 confocal Z-planes. Scale bar = 10 μm.

**S9 Movie. Cortical actomyosin inhibition (Cdc42T17N).**

Live fluorescence and DIC imaging of activated oocytes injected with the H2b-mCherry DNA marker alone (control, left panel) or a combination of H2b-mCherry and a Cdc42 dominant negative form (Cdc42T17N, small polar body (middle) and spindle relocation (right) phenotype). The oocyte have been injected with the H2b-mCherry DNA marker. Images were acquired every 2 min with a 20x objective and represent a maximum projection of 20 confocal Z-planes (H2b-mCherry) or a single confocal Z-plane (DIC). Scale bar = 10 μm.

**S10 Movie. Cortical actomyosin inhibition (Cdc42T17N) (PIV).**

Live fluorescence and DIC imaging of activated oocytes injected with the H2b-mCherry DNA marker alone (control, left panel) or a combination of H2b-mCherry and a Cdc42 dominant negative form (Cdc42T17N, small polar body (middle) and spindle relocation (right) phenotype). The oocyte have been injected with the H2b-mCherry DNA marker. Images were acquired every 2 min with a 20x objective and represent a maximum projection of 20 confocal Z-planes (H2b-mCherry) or a single confocal Z-plane (DIC). The PIV vector field enables to visualize cytoplasmic streaming and the color code indicates the speed of tracked particles. Scale bar = 10 μm.

**S11 Movie. Antagonistic forces yield symmetry breaking and spindle rotation.**

Time-lapse sequences of typical simulations with (left) or without (middle and right) Gaussian noise added to the DNA clusters position.

## Materials and methods

### Oocyte recovery and culture

All animal procedures were conducted in accordance with the European directive for the use and care of laboratory animals (2010/63/EU), and approved by the local animal ethics committee under the French Ministry of Higher Education, Research and Innovation (Project licence APAFIS#11761). Female mice of the OF1 strain (8-10 week old; Charles River) were primed by intraperitoneal injection of 5-7 units of PMSG (Chronogest, MSD), followed 48h later by a second injection of 5 IU hCG (Chorulon, MSD). Metaphase-II (MII) oocytes were recovered from the oviducts in M2 medium (Sigma) supplemented with 3mg/ml hyaluronidase (Sigma). To induce resumption of meiosis-II, MII oocytes were treated for 7 minutes with 7% ethanol, as previously described (23). When indicated, the culture medium was supplemented with the PLK1 inhibitor Bi-2536 (250 nM; Axon Medchem #1129) or the Rock-1 inhibitor Y-27632 (50 μM; Merck Millipore #688000). For activated oocytes experiments in Fig 2C-G and S3B Fig, Bi-2536 and Y-27632 were added to the culture medium immediately after the activation. For metaphase II experiments in S3A Fig, the oocytes have been incubated with Y-27632 for 2 hours before fixation.

### Plasmids, cRNA preparation and microinjection

The following plasmids were used: H2B-mCherry in pcDNA3 (Robert Benezra; Addgene plasmid #20972; (51)), pGEMHE-eGFP-MAP4 (Jan Ellenberg; Euroscarf plasmid #P30518; (52)), eGFP-UtrCH in pCS2+ (William Bement; Addgene plasmid #26737; (46)), pRK5-myc-Cdc42-T17N (Gary Bokoch; Addgene plasmid #12973), which we subcloned into pcDNA3.1 and RanT24N in pcDNA3.1 (Ben Margolis; (53)). The full-length Ect2 sequence was amplified from mouse oocyte cDNA and cloned into pcDNA3.1 downstream of eGFP, to generate the eGFP-Ect2 construct. Polyadenylated cRNAs were synthesized in vitro from linearized plasmids, using the mMessage mMachine T7 or SP6 kit and Poly(A) Tailing kit (Ambion), purified with RNeasy purification kit (Qiagen), and stored at −80°C. Using the electrical-assisted microinjection technique, which allows for a high rate of oocyte survival (54), MII oocytes were injected with ~5pl cRNA and cultured for ~3 hours to allow for protein expression.

### Immunofluorescence

For actomyosin staining, the oocytes were fixed for 25 min at room temperature (RT) with paraformaldehyde 3% in PBS, freshly prepared from a 16% methanol-free paraformaldehyde solution (Electron Microscopy Sciences) and adjusted to pH ~7.5 with NaOH. For RhoA, Anillin and Rock-1 staining, oocytes were fixed for 10 min at 4°C with pre-cooled 10% trichloroacetic acid (Sigma). Next, fixed oocytes were then permeabilized with 0.01% Triton X-100 (Sigma) in PBS for 15 minutes at RT, blocked with 3% BSA (Sigma) in PBS for 2 h at RT, and incubated overnight at 4°C with primary antibodies in PBS-BSA 3%. On the next day, oocytes were washed in PBS-BSA 3% and incubated with secondary antibodies diluted 1:1000 in PBS-BSA 1%, for 45 min at 37°C. The following primary antibodies were used: phospho-myosin light chain 2 (Ser19), pMLC2 (1/200 dilution, Rabbit polyclonal, Cell Signaling Technology #3671), RhoA (1/100 dilution, mouse monoclonal; Santa Cruz sc-418), Anillin (1/100, Rabbit polyclonal; Santa Cruz sc-67327), Rock-1 (1/100, Goat polyclonal; Santa Cruz sc-6056). Secondary antibodies were Alexa Fluor 488-conjugated donkey anti-goat, donkey anti-rabbit and goat anti-mouse (Invitrogen). Actin filaments were stained with Alexa Fluor 568-phalloidin (1/50 dilution, 66 μM, Life Technologies). Chromatin was stained with To-Pro-3 (1/200 dilution, 1 mM, Invitrogen T3605).

### Live microscopy

Oocytes were placed on glass-bottom dishes (MatTek, Ashland, MA) in a small drop of medium covered with mineral oil, and imaged with a Leica SP5 or SP8 confocal microscope, using a 63x or 20x oil-immersion objective. For 63x live imaging, a single confocal Z-plane was acquired every 1 min. For 20x live imaging, 20 confocal Z-planes separated by 3.2 μm were acquired every 2 min. In both cases, the temperature was maintained at 37°C using a stage top incubator (INUBG2E-GSI, Tokai Hit, Shizuoka-ken, Japan) fitted on the microscope stage.

### Image processing and data analysis

#### Software

All image processing and data analyses were performed using ImageJ 1.52t or Matlab 2015a either separately or together using the MIJ plugin (D. Sage, D. Prodanov, C. Ortiz and J. Y. Tivenez, retrieved from http://bigwww.epfl.ch/sage/soft/mij). The graphics were produced using OriginPro 9.0 software and exported to Adobe Illustrator CS6 for final processing.

#### Live 3D segmentation and tracking procedure

To monitor the position of the chromatids clusters within the shell of the oocyte, we developed an automatic 3D segmentation and tracking procedure coded in Matlab and ImageJ. This method, based on grey levels thresholding, enabled us to extract the DNA clusters and the cell outlines in H2b-mCherry injected oocytes. Briefly, we filtered the images using Gaussian blurs to smoothen the edges and applied adaptive thresholding to account for the overall brightness reduction in the deeper part of the Z-stacks. We then used different threshold parameters to separate the DNA clusters from the cytoplasmic signal and finally tracked the DNA clusters over time using a proximity criterion. This method allowed us to extract the following parameters: <u>The distance between the DNA clusters</u> which we obtained by measuring the 3D Euclidian distance between the two DNA clusters. We used the variation of this distance to register in time the oocyte populations. To do so, we manually selected early time-points for which the distance increases linearly. We next fitted these first few time-points and defined the intercept of the linear fit and the time axis as t=0 (see Fig 1E). <u>The spindle rotation angle</u> which we obtained by measuring in 3D the angle formed by the internalized cluster of chromatids (C_In_), the spindle mid-point (equidistant from the two clusters) and the oocyte centroid (see Fig 1F). <u>The DNA clusters distance to cortex</u> which we obtained by averaging the 3D Euclidian distance between the DNA clusters and their 1% closest cortical pixels, resulting from the cytoplasmic outlines segmentation. <u>The spindle mid-point distance to the oocyte centroid</u> which we obtained by measuring the 3D Euclidian distance between the spindle mid-point (equidistant from the two clusters) and the oocyte centroid. <u>The initial time of rotation (t_i_ rotation)</u> which we defined as the time when the spindle reaches 5% of its total fitted rotation. To fit the rotation, we manually selected the time-points corresponding to the actual rotation and fitted the results using a 5 parameters logistic model implemented in Matlab (G. Cardillo, retrieved from https://github.com/dnafinder/logistic5).

#### Cortical F-actin and myosin-II levels and distance to DNA clusters

During this study we repeatedly assessed the cortical F-actin and myosin-II levels and eventually correlated them to the distance existing between the cortex and the DNA clusters (see Fig 3B, 3C, 4B, 4C and 5C). To do so, we first segmented the oocyte, using the procedure described above, and defined a 10 to 20 pixels wide line selections following the cell outline. We next generated Euclidian distance maps (see EDM commands in both Matlab and ImageJ) centered on the centroid of the DNA clusters. These maps use grayscales to encode the distances between a previously defined point (e.g. DNA cluster centroid) and the rest of the image pixels. Finally, we extracted the values along the line selections for the F-actin, myosin-II and EDM channels. In Fig 3B (cortical line-scans), we simply plotted the cortical F-actin and myosin-II levels as a function of the linear cortical distance to the cortical point situated at equal distance from the two DNA clusters. In Fig 3C and 5C, we averaged profiles from different oocytes showing the cortical F-actin and myosin-II levels as a function of the Euclidian distance from the cortex to the DNA clusters. Fig 4B, we generated 2D color-coded maps of the polarized domain showing the variation over time of either the cortical F-actin levels or the cortical distance to the DNA clusters. In Fig 4C, we recapitulated the color-coded maps in 1D by averaging results for the 25% (cortical F-actin levels) or the 1% (cortical distance to the DNA clusters) closest cortical pixels.

#### Particle image velocimetry (PIV)

To measure the flow of cytoplasmic materials in activated oocytes, we devised a Matlab routine built on the PIVlab implementation of particle image velocimetry (55). To avoid any interference with the analysis, we eliminated the vectors outside the oocyte using a mask. We smoothed the vector field in space and time with custom filters based on running averages. Finally, for display purposes, we color-coded the vector norm using the “quiverc” function (B. Dano, retrieved from https://www.mathworks.com/matlabcentral/fileexchange/3225-quiverc). In S1A Fig, we used an interrogation window of 80 pixels for the first pass and 40 pixels for the second pass. We smoothed data using a 5×5 vectors kernel in space and a 9 time-points kernel in time. In S1B Fig, Fig 2G and 5G, we used an interrogation window of 80 pixels for the first pass and 40 pixels for the second pass. We smoothed data using a 3×3 vectors kernel in space and a 7 time-points kernel in time.

### Numerical modelling

In the model, we considered a spindle of fixed length in a circular oocyte. DNA clusters are localized at the ends of the spindle. As described in the main text, the spindle is essentially submitted to two opposing forces: the attraction of each cluster to the locally polarized cortex, and the cytokinetic ingression pushing the central spindle inward. A schematic of the model geometry is depicted Fig 6A.

### Attraction to the cortex

We modeled the attraction of DNA clusters to the cortex through an effective potential. For simplicity, we use a soft-core-like potential *V* of the form (Fig 6B):

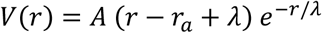

*r* is the distance to the attraction zone of the cortex, *λ* is the typical decay length of the potential. *r_a_* is the value where *V*(*r*) is minimum, and typically corresponds to the radius of DNA clusters: *V* is minimum when the cluster is apposed to the attraction zone.

The force 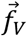 exerted by each attraction zone on the corresponding cluster is simply the gradient of *V*(*r*):

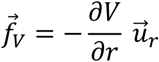

### Central ingression

The cytokinetic ingression acts as a pushing force on the central spindle. For simplicity, we assume in the model that the ingression velocity is constant. Numerically, we simply prescribe the velocity of the central spindle, which moves with constant velocity 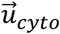 perpendicular to the spindle.

Importantly, without this prescribed ingression, the clusters remain at the cortex and nothing happens. This corresponds to the arrested state of the oocyte.

### Spindle flexibility

We allow the spindle to slightly bend. Bending of the spindle yields an elastic force on the clusters, that is proportional to the deflection 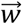 of the spindle and to the spindle stiffness *k*, so that the elastic force is 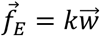. Note that 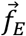 is perpendicular to the axis joining the clusters.

### Time evolution

We obtain the movement of each cluster using overdamped dynamics:

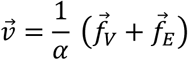

Where 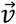 is the velocity of a given cluster, and *α* the damping coefficient. To compute the new position of each cluster at each time step, we integrate this equation numerically using a simple Euler method.

### Noise

This problem has an unstable solution, in which the spindle remains parallel to the cortex while it is pushed by the ingression. In that case, both clusters eventually escape their attraction potential. This unstable solution cannot be observed in vivo, as any noise, asymmetry or perturbation will lead to symmetry breaking. The unstable solution can be observed numerically if no noise whatsoever is introduced (Fig 6C), as the simulation is then perfectly symmetric. However, as soon as a very small noise is introduced, the system eventually breaks its left-right symmetry, with one cluster escaping its attraction potential while the other returns to its attraction zone (Fig 6C).

There are many possible sources of noise in the biological system, which may include (but are not limited to): an asymmetry of the initial configuration, an asymmetry of the decay length *λ*, an symmetry of the attraction fo thermal or active fluctuations of the clusters, an imperfect centering of the central spindle or of the ingression, etc. In this article, we only consider noise on the cluster’s dynamics.

Upon integration of the equations of motion, we add a normally distributed random noise on each cluster’s position at each time step, with standard deviation *σ*. This is sufficient to break the symmetry of the system even when *σ* is very small compared to the system size. Note that this also introduces a normally distributed initial asymmetry, which of course sets a bias for the direction of rotation (Fig 6F).

### Parameters values

In the simulations, we mostly use the same set of parameters throughout the paper. The only changes concern the noise introduced to make better sense of the symmetry-breaking mechanism and of the variability observed experimentally. Note that distances are in units of *R_ovo_*, the radius of the oocyte (we set *R_ovo_* = 1). The timescale is arbitrary, as we set the damping coefficient to 1.

Parameters used throughout the paper:

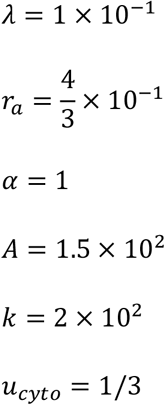

Noise:

Fig 6C (simulations, top panel): *σ* = 0

Fig 6C (simulations, other panels): *σ* = 1 × 10^−3^

Fig 6D, 6E and 6F (angles & distances dynamic): *σ* = 1 × 10^−3^

Fig 6G (int/out histogram, in/out & rotation time): *σ* = 1 × 10^−2^

### Simplified analytical model of the “symmetry-breaking time” distribution

To make better sense of the long-tailed distribution of the symmetry breaking time, observed both in vivo and in silico, we designed a simplified model of the symmetry-breaking process. Let us consider the dynamics of an object on a locally quadratic bump of potential *V*:

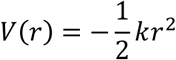

where *r* is the position of the object and *a* is a constant. The overdamped dynamics of the object is simply given by:

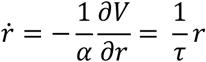

where *α* is the damping coefficient, and 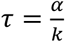 is the relaxation timescale. The position of the object at time *t* is thus given by:

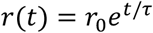

If *r*_0_ = *r*(*t* = 0) = 0, the solution is *r*(*t*) = 0 (the object remains on the maximum of *V*(*r*). Clearly this is an unstable solution, as any infinitesimal deviation from *r*_0_ = 0 will increase exponentially over time, with the object moving away from the maximum of *V*(*r*). Let us now consider the time *t*_*c*_ (the “symmetry-breaking” time) needed for the object to reach an arbitrary distance *r*_*c*_ considered the symmetry-breaking distance, starting from *r*_0_ ≠ 0. Taking the log of the equation above, we have:

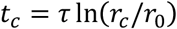

Let us now assume that *r*_0_ has a Gaussian probability density function (with standard deviation *σ*), as in our batch of oocytes simulations: 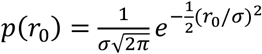. Using the expression of *t*_*c*_ above and the probability density function of *r*_0_, one can directly calculate the probability density function of the symmetry-breaking time *t*_*c*_, which reads:

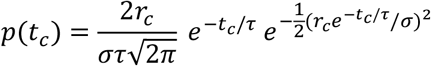

With the analytical expression above we directly recover the long-tailed distribution *p*(*t*_*c*_) observed in our simulations and in vivo, starting from a Gaussian noise (S4 Fig, top and middle panels). Note that computing *p*(*t*_*c*_) numerically from a normal distribution of *r*_0_ yields the same result. The long tail of *p*(*t*_*c*_) can be viewed as a direct consequence of the non-linear decay of *t*_*c*_(*r*_0_) close to the unstable solution. Small values of *r*_0_ lead to a long symmetry-breaking time (Figure S4 Fig, bottom panel). For plots shown Figure S4 Fig, we used *σ* = 1, *τ* = 1, and *r*_*c*_ = 10.

### Statistics and reproducibility

Data points from different oocytes were pooled to estimate the mean and standard deviations. The quantifications were carried out on a minimum number of 10 oocytes from a minimum number of two independent experiments (mounting). Statistical significance was tested using Mann–Whitney tests, assuming non-normal distributions and equal variances. For the data in Fig 6G (top row), we performed a one-way ANOVA to test the influence of the Cortex/DNA clusters distance ratio over the initial time of rotation (t_i_ rotation). No statistical method was used to predetermine sample sizes. The experiments were not randomized, and the investigators were not blinded to allocation during experiments and outcome assessment.

